# Nationwide Spread of Fluconazole-Resistant *Candida parapsilosis* Clones: Insights from the Antifungal Resistance Surveillance Program

**DOI:** 10.64898/2026.08.10.740302

**Authors:** Elena López-Peralta, Cristina de Armentia-Roldán, Alejandra Roldán, Susana Sánchez-Galiano, Maite Ruiz Pérez de Pipaón, Irene Merino, Marta López-Lomba, María Teresa Durán-Valle, Paloma Merino-Amador, Fernando González-Romo, María Teresa Martín-Gómez, Mireia Puig-Asensio, Carmen Ardanuy, Julio García-Rodríguez, Alfredo Maldonado-Barrueco, Gregoria Megías-Lobón, María Ángeles Mantecón-Vallejo, María Antonia Miguel Gómez, Teresa Nebreda-Mayoral, Octavio Carretero Vicario, Mercedes Delgado-Valverde, Inés Portillo-Calderón, Natalia Chueca-Porcuna, Mónica Chávez-Caballero, Concepción Mediavilla-Gradolph, María Evangelina Pablo Hernando, Marta Arias-Temprano, María Pía Roiz-Mesones, Isabel Lara-Plaza, Leyre López-Soria, José Luis Barrios-Andrés, Clara Lejarraga-Cañas, Mateu Espasa, Antonio Casabella-Pernás, Ana Pérez Ayala, Isabel Sánchez-Romero, María Muñoz-Algarra, Sarela García Masedo-Fernández, Albert Bas-Vilda, Elena Cuadros-Moronta, Carla Iglesias-Escobar, Eva Alcoceba, Celia Garcia-Rivera, Juan Cuadros, Elía Gómez G. de la Pedrosa, Manuel Monsonis, Cristina Esteva, Eneritz Velasco, Julia Gotzens, Sarai Varona, Sara Monzón, Isabel Cuesta, Miquel Àngel Schikora-Tamarit, Toni Gabaldón, Óscar Zaragoza, Laura Alcázar-Fuoli

**Affiliations:** Mycology Reference Laboratory. National Centre for Microbiology. Instituto de Salud Carlos III, Madrid, Spain; Clinical Unit of Infectious Diseases, Microbiology and Parasitology. Virgen del Rocío Universitary Hospital. Seville, Spain. Clinical and Molecular Microbiology Group. Institute of Biomedicine of Seville. HUVR/CSIC/University of Seville.; Center for Biomedical Research Network in Infectious Diseases (CIBERINFEC; CB21/13/00006), Instituto de Salud Carlos III, Madrid, Spain; Microbiology Unit. Universitary Hospital Rio Hortega. Valladolid, Spain; Microbiology and Parasitology Department. Móstoles Universitary Hospital. Madrid, Spain; Microbiology Department, Universitary Hospital Clínico San Carlos, Madrid, Spain; Health Research Institute from Hospital Clínico San Carlos (IdISSC), Madrid, Spain; Department of Medicine, Complutense University, School of Medicine, Madrid, Spain; Department of Microbiology, Vall d’Hebron University Hospital, Universitat Autònoma de Barcelona, Barcelona 08035, Cataluña, Spain; Department of Infectious Diseases, Bellvitge Universitary Hospital. Biomedical Research Institute of Bellvitge Institut d’Investigació Biomèdica de Bellvitge, IDIBELL), Barcelona, Spain; Center for Biomedical Research Network in Infectious Diseases (CIBERINFEC; CB21/13/00009), Instituto de Salud Carlos III, Madrid, Spain; Microbiology Department, Bellvitge Universitary Hospital. Biomedical Research Institute of Bellvitge Institut d’Investigació Biomèdica de Bellvitge, IDIBELL), Barcelona, Spain; Center for Biomedical Research Network in Respiratory Diseases Enfermedades (CIBERES CB06/06/0037), Instituto de Salud Carlos III, Madrid, Spain; Clinical Microbiology and Parasitology Department, Hospital Universitario La Paz, IdiPaz, Madrid, Spain; Center for Biomedical Research Network in Infectious Diseases (CIBERINFEC; CB21/13/00039), Instituto de Salud Carlos III, Madrid, Spain; Department of Clinical Microbiology. Burgos Universitary Hospital. Castilla y León. Spain; Microbiology and Immunology Unit. Valladolid Clinical Universitary Hospital, Castilla y León, Spain; Microbiology and Parasitology Unit. Salamanca Universitary Hospital. Castilla y León, Spain; Microbiology Service, Clinic Unit of Infectious Diseases and Microbiology, Hospital Universitario Virgen Macarena, Seville, Spain. Antimicrobial resistance and complex infectious group. Instituto de Biomedicina de Sevilla, Hospital Universitario Virgen Macarena/CSIC/Universidad de Sevilla, Sevilla, Spain; Center for Biomedical Research in Network in Infectious Diseases (CIBERINFEC-CB21/13/00012), Instituto de Salud Carlos III, Madrid, Spain; Service of Microbiology, Hospital Universitario San Cecilio, Granada, Spain. Instituto Biosanitario de Granada (ibs.GRANADA), Granada, Spain; Center for Biomedical Research in Network in Infectious Diseases (CIBERINFEC-CB21/13/00088), Instituto de Salud Carlos III, Madrid, Spain; Microbiology Unit. San Juan de Dios del Aljarafe Hospital, Sevilla, España; Microbiology Unit. Malaga Universitary Hospital. Andalusia, Spain; Hospital Clínico Universitario Lozano Blesa, Avda. San Juan Bosco 15, 50009, Zaragoza, Spain; Hospital de Cabueñes, Asturias, Spain; Microbiology Department, Marqués de Valdecilla Universitary Hospital, Santander. Cantabria. Spain. Valdecilla Research Instituto (Instituto de Investigación Valdecilla, IDIVAL). Spain; Center for Biomedical Research in Network in Infectious Diseases CIBERINFEC (CB21/13/00068), Instituto de Salud Carlos III, Madrid, Spain; Microbiology Service. Hospital Universitario Cruces, Barakaldo, Vizcaia, Spain; Servei de Microbiologia, Hospital Universitari Parc Taulí, Institut d’Investigació i Innovació Parc Taulí (I3PT-CERCA), Universitat Autònoma de Barcelona, Sabadell, Spain; Microbiology Unit. Universitary Hospital 12 de Octubre. Madrid, Spain. Research Institute from Hospital 12 de Octubre (i + 12 Institute), Madrid, Spain; Microbiology Department, Puerta de Hierro Universitary Hospital, Majadahonda, Madrid, Spain; Department of Microbiology and Parasitology. Hospital Mateu Orfila. Maó, Menorca, Balearic Islands, Spain; Microbiology Unit, Hospital Universitary Son Espases, Palma de Mallorca, Spain. Research Group on antibiotic resistance and bacterial infections pathogenesis, Health Research Institute of the Balearic Islands (IdISBa).; Microbiology Department. General Universitary Hospital Virgen de los Lirios. Alcoy, Alicante, Spain; Microbiology Department, Hospital Príncipe de Asturias, Madrid, Spain; Microbiology Department, Ramón y Cajal University Hospital, IRYCIS; Center for Biomedical Research in Network in Infectious Diseases CIBERINFEC (CB21/13/00084), Instituto de Salud Carlos III, Madrid, Spain; Microbiology Service. Hospital Sant Joan de Deu. Barcelona, Spain; Infectious Diseases and Systemic Inflammatory Response in Pediatrics, Pediatric Infectious Diseases Department, Institut de Recerca Sant Joan de Déu, Hospital Sant Joan de Déu, Barcelona, Spain; Bioinformatics Unit, Instituto de Salud Carlos III, Madrid, Spain; Barcelona Supercomputing Centre (BSC-CNS), Barcelona, Spain; Institute for Research in Biomedicine (IRB Barcelona), The Barcelona Institute of Science and Technology, Barcelona, Spain; Center for Biomedical Research in Network in Infectious Diseases CIBERINFEC (CB21/13/00105), Instituto de Salud Carlos III, Madrid, Spain; Catalan Institution for Research and Advanced Studies (ICREA), Barcelona, Spain

**Keywords:** *Candida parapsilosis*, fluconazole, Erg11, antifungal resistance, antifungal susceptibility testing

## Abstract

**Background:** Outbreaks of fluconazole-resistant *Candida parapsilosis* have recently emerged worldwide. In Spain, this phenomenon has been reported since 2020, mainly involving isolates from different clones harbouring the Y132F mutation at Erg11.

**Methods:** We analysed the expansion of fluconazole resistant *C. parapsilosis* strains within the national antifungal resistance surveillance program. Genetic clustering and relationships were assessed using microsatellite typing and whole genome sequencing.

**Findings:** We identified the expansion of three distinct clones carrying the Y132F mutation. Additionally, there was an increase in strains harbouring the G458S mutation, most of which belonged to a clonal complex, although other less prevalent clones were also detected. G458S isolates showed higher resistance to azoles than Y132F strains, particularly to voriconazole and isavuconazole. This increased resistance was associated with mutations in the Tac1 transcriptional regulator and duplication of a chromosomal region containing Tac1 and Erg11. One G458S isolate without mutation at Tac1 exhibited lower MIC values. Furthermore, two isolates carried the K143R mutation, and a distinct group of resistant strains without detectable *ERG11* mutations was also identified. Resistant cases were detected across 31 hospitals in 12 autonomous regions.

**Interpretation:** Our findings indicate a concerning nationwide expansion of antifungal-resistant *C. parapsilosis* in Spain, involving multiple resistance mechanisms and clonal lineages, with implications for antifungal treatment and infection control strategies.

## Introduction

Fluconazole resistance in *Candida parapsilosis* is one of the main epidemiological concerns related to antifungal resistance that has emerged in the last years. Globally, this yeast ranks as the second or third most common cause of bloodstream yeast infections after *Candida albicans* and *Nakaseomyces glabratus* (formerly *Candida glabrata*) [1–4].

*Candida parapsilosis* infections are particularly prevalent among neonates, immunocompromised individuals, and patients with central venous catheters or parenteral nutrition [1, 5]. Furthermore, *C. parapsilosis* is associated to nosocomial outbreaks and transmission through direct and indirect contact, either via the hands of healthcare workers or through contaminated patient care equipment [6, 7].

Acquired resistance to fluconazole in *C. parapsilosis* has been recently observed worldwide, often in the context of nosocomial outbreaks [8–30]. In Spain, different studies have reported a significant rise, since 2020, in the number of fluconazole non-susceptible (*FNS*) *C. parapsilosis* in various tertiary hospitals [31–33].

Several molecular mechanisms involved in fluconazole resistance have been identified, including point mutations in Erg11 (14-sterol-α-demethylase, target of triazoles), overexpression of the *ERG11* gene, and mutations in other *ERG* genes [8, 34]. The Y132F substitution at Erg11 is the dominant amino acid change associated with resistance to both fluconazole and voriconazole. Other Erg11 substitutions linked to fluconazole resistance, though less commonly reported, include G458S, D421N, K128N, and K143R [10, 21, 32, 35–37]. Overexpression of MFS transporters (such as *MDR1*) and ABC efflux pumps (*CDR* genes), and mutations at transcription factors involved in the regulation of expression of these transporters (such as Mrr1 and Tac1) are also related to fluconazole resistance [38–43].

In this study, we present updated data resistance to fluconazole of *C. parapsilosis* sensu stricto reported to the Mycology Reference Laboratory of National Centre for Microbiology from January 2022 to December 2025. We report an increase and expansion of strains carrying the Erg11^G458S^ mutation. The majority of these strains also exhibited mutations in the Tac1 transcription factor. Additionally, we describe the continued geographical dissemination of clones harbouring the Erg11^Y132F^ mutation.

## Material and Methods

### Media and strains identification

The study was performed on 1,107 clinical isolates received in the National Centre for Microbiology from January 2022 to December 2025. This study included all resistant strains from the studied period and susceptible strains isolated from hospitals where at least one resistant strain was reported. Species identification was confirmed by sequencing the ITS region from the ribosomal DNA [44].

### Antifungal susceptibility

Antifungal susceptibility was confirmed at the Mycology Reference Laboratory (Instituto de Salud Carlos III) using the EUCAST protocol (https://www.eucast.org/fileadmin/eucast/pdf/AFST/methodology/EUCAST_E.Def_7.4_Yeast_definitive_revised_2023.pdf). Briefly, RPMI 1640 medium (Merck, Sigma-Aldrich) was buffered with MOPS (Merck, Sigma-Aldrich) at pH 7 and supplemented with 2% glucose (Merck, Sigma-Aldrich). The following antifungal concentration ranges were used: Amphotericin B (AmB, Merck, Sigma-Aldrich, 16-0.03 mg/L), flucytosine (64-0.125 mg/L), fluconazole (FLC, Merck, Sigma-Aldrich, 64-0.125 mg/L), itraconazole (ITZ, Janssen Pharmaceutical Research and Development, 8-0.016 mg/L), voriconazole (VOR, Pfizer Pharmaceutical Group, 8-0.016 mg/L), posaconazole (POS, Merck, Sigma-Aldrich, 8-0.016 mg/L), isavuconazole (ISV, Pfizer Pharmaceutical Group, 8-0.016 mg/L), caspofungin (CSP, Merck, Sigma-Aldrich, 16-0.016 mg/L), micafungin (MICA, Astellas Pharma Inc, 2-0.004 mg/L) and anidulafungin (ANID, Pfizer Pharmaceutical Group, 4-0.008 mg/L). The minimal inhibitory concentration (MIC) was defined as the concentration that inhibited growth by 50% related to the control without antifungal, except for AmB, when 90% inhibition was considered. Susceptible (S), resistant (R) or susceptible increased exposure (I) strains were defined as established by EUCAST breakpoints v.12.0 (https://www.eucast.org/fungi-afst/clinical-breakpoints-and-interpretation/clinical-breakpoint-table/, June 26^th^, 2025). As quality control strains, we included *C. parapsilosis* ATCC 22019 and *C. krusei* ATCC 6258 isolates.

### Typing by microsatellites analysis

Typing of the *C. parapsilosis* strains was performed by analysis of microsatellites regions as described in [45], with the modifications described in [31]. As internal size standard, GeneScan ROX 500 was added to all the samples. The PCR products were separated in a AB3730XL DNA analyzer (Applied Biosystems), and sizes analysis was performed with Peak Scanner V 2.0 software (Applied Biosystems).

Similarities between genotypes were visualized by constructing a minimum spanning tree using Bionumerics, version 8.1 (Applied Maths, St.-Martens-Latem, Belgium), treating the data as categorical information.

Genotypes were assigned based on the number of repetitions in each marker. Strains were considered from the same genotype when the strains showed the same repetitions. Strains that only showed a difference in one marker were considered clonal complexes (also referred as clones) [45].

### Identification of mutations at the *ERG11* gene

*ERG11* gene was amplified and sequenced as described in [31] with the modification described in [46]. Sequences were analyzed with Seqman software (DNA Lasergene 12 package) to identify mutations in *ERG11* gene compared to the sequence from the QC strain ATCC 22019 (GenBank accession no. GQ302972). Alternatively, *ERG11* mutations were also detected using a real time PCR method described in [46]. This last method was mainly applied to strains belonging to the main clones harboring *ERG11* mutations.

### Whole genome sequencing of *C. parapsilosis* strains

We obtained whole genome sequences from [47] and available in SRA through PRJNA1439216, which included representative isolates from different genotypes and different resistant mechanisms. Moreover, we sequenced the genomes of additional strains 24 strains (PRJNA1476012) that increased the representative of strains obtained from different hospitals. Briefly, DNA was obtained using the phenol-chloroform method after disrupting the cells with glass beads using FastPrep (MP Biomedicals). Libraries were prepared using DNA Prep. Tagmentation kit (Illumina) following the manufactureŕs protocol. Sequencing was done in a NovaSeq sequencer (NovaSeq 6000 SP Reagent kit v1.5, double stranded).

### Bioinformatic analysis

Whole-genome sequencing reads were analysed using the nf-core/sarek v3.1.1 pipeline (https://github.com/nf-core/sarek) [48] implemented in Nextflow by the nf-core community [49]. The workflow started with quality control of raw reads using FastQC v0.11.9 (https://www.bioinformatics.babraham.ac.uk/projects/fastqc/), followed by read trimming and filtering with fastp v0.23.2 [50]. Trimmed reads were mapped against the reference genome of *Candida parapsilosis* strain CDC317 (assembly accession GCF_000182765.1, ASM18276v2) using BWA-MEM v0.7.17-r118 [51]. Variant calling was performed using FreeBayes v1.3.6 [52] and MultiQC v1.13 [53] was used to aggregate metrics across the pipeline results.

A SNP alignment was generated from the merged variant calling format (VCF) file using the generate_snp_sequence.R script distributed with SNPhylo v 20180901 (https://github.com/thlee/SNPhylo). SNP filtering parameters included a linkage disequilibrium threshold of 2 and a minimum allele frequency threshold of 0.5. Maximum likelihood phylogenetic inference was performed using IQ-TREE v2.2.6 [54]. ModelFinder Plus (MFP) was first used to identify the best-fitting nucleotide substitution model for the SNP alignment. The best-fitting model identified was GTR+F+R2. Final phylogenetic reconstruction was then performed using this model with 1,000 ultrafast bootstrap replicates. The final tree display was generated using v.7 (https://itol.embl.de/).

Aligned documents (.bam) were visualized with the software Integrated Genomic Viewer (IGV, version 2.8.13) using the genome of the CDC317 strain as reference. This genome has been largely used as reference in many genomic analysis, and it contains the Y132F mutation at Erg11 in heterozygosis. Global coverage was estimated using the “Count” option from igvtools, using an average window of 500 bp.

### Detection of the number of copies of *ERG11* by real time PCR

To quantify the number of copies of ERG11 compared to ACT1 (CPAR2_201570, unicopy control), both genes were amplified by real-time PCR using the oligonucleotides described in **Table 1**. 200 pg of DNA were used as template in a reaction containing 800 nM of each oligonucleotide, 750 µM of MgCl_2_ and 10 µL of 2x SensiMix SYBR Hi-ROX Kit (total final volume of 20 µL). Amplification was performed using the following conditions: 10 minutes at 95 °C, 50 cycles of 10 seconds at 95 °C, 5 seconds at 55 °C and 30 seconds at 72 °C, with a final cycle of 5 seconds at 65 °C, 1 minute at 97 °C and 30 seconds at 40 °C. Then, the crossing threshold value (Ct) for each reaction was calculated using the software. Estimated variation in the number of copies of *ERG11* in a strain (StX) compared to the control strain (StC) normalized by the copies of *ACT1* was estimated using the following formula:

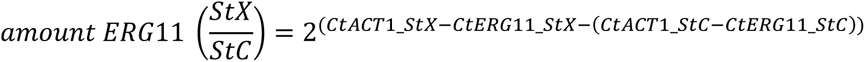

**Table 1:**
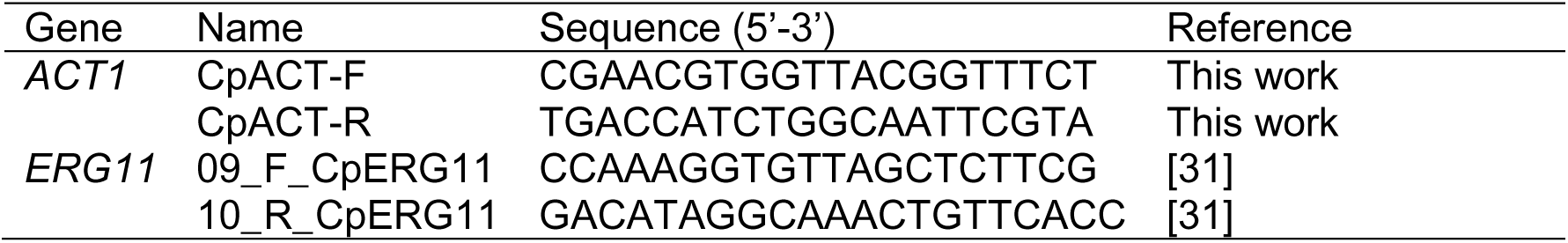
Oligonucleotides used to detect the *ERG11* copy number in *C. parapsilosis*.

### Statistics

MIC data was analysed using Excel (Microsoft office 365) and GraphPad Prism 9 softwares. To evaluate statistical differences in antifungal susceptibility, MIC values were log(2) converted, and differences between the groups were assessed with ANOVA test using Tukey post-hoc test for multiple comparisons. Differences were considered significant when p-value < 0.05.

## Results

Resistant strains were collected from 31 hospitals across 12 autonomous regions (17 provinces) in Spain (**Figure 1**). A summary of the number and geographical distribution of the strains is presented in **Table 2**. According to the site of isolation, strains were classified as environmental, urine, colonization, or clinical samples from superficial or invasive infections (**Table 3**).

**Figure 1:**
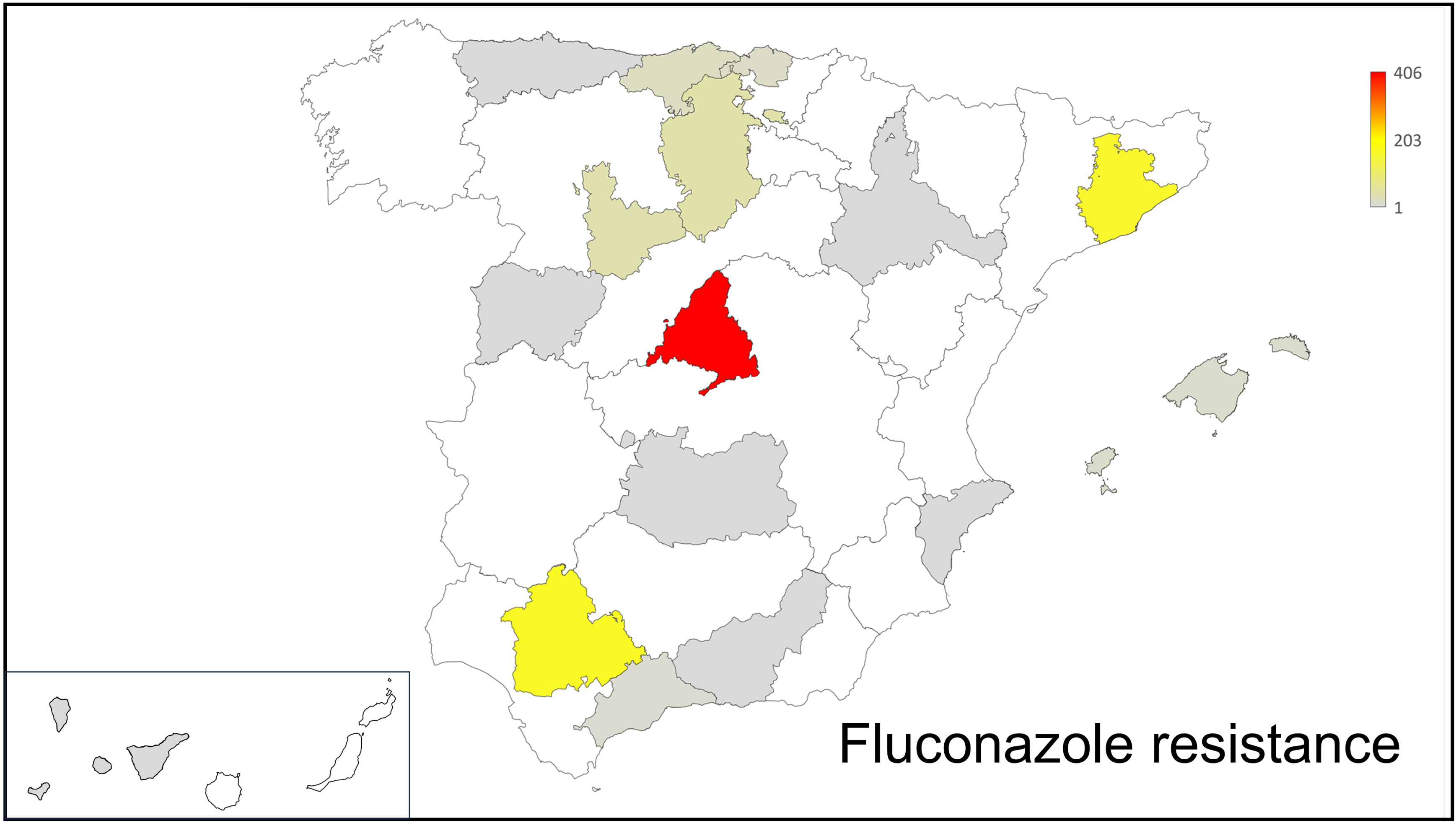
Global distribution of fluconazole-resistant strains in Spain, shown as a choropleth map in which each province is colored according to the total number of resistant strains. The color scale ranges from light grey (minimum value, 1) to deep red (maximum value, 406), with intermediate values represented by a continuous red gradient.

**Table 2:**
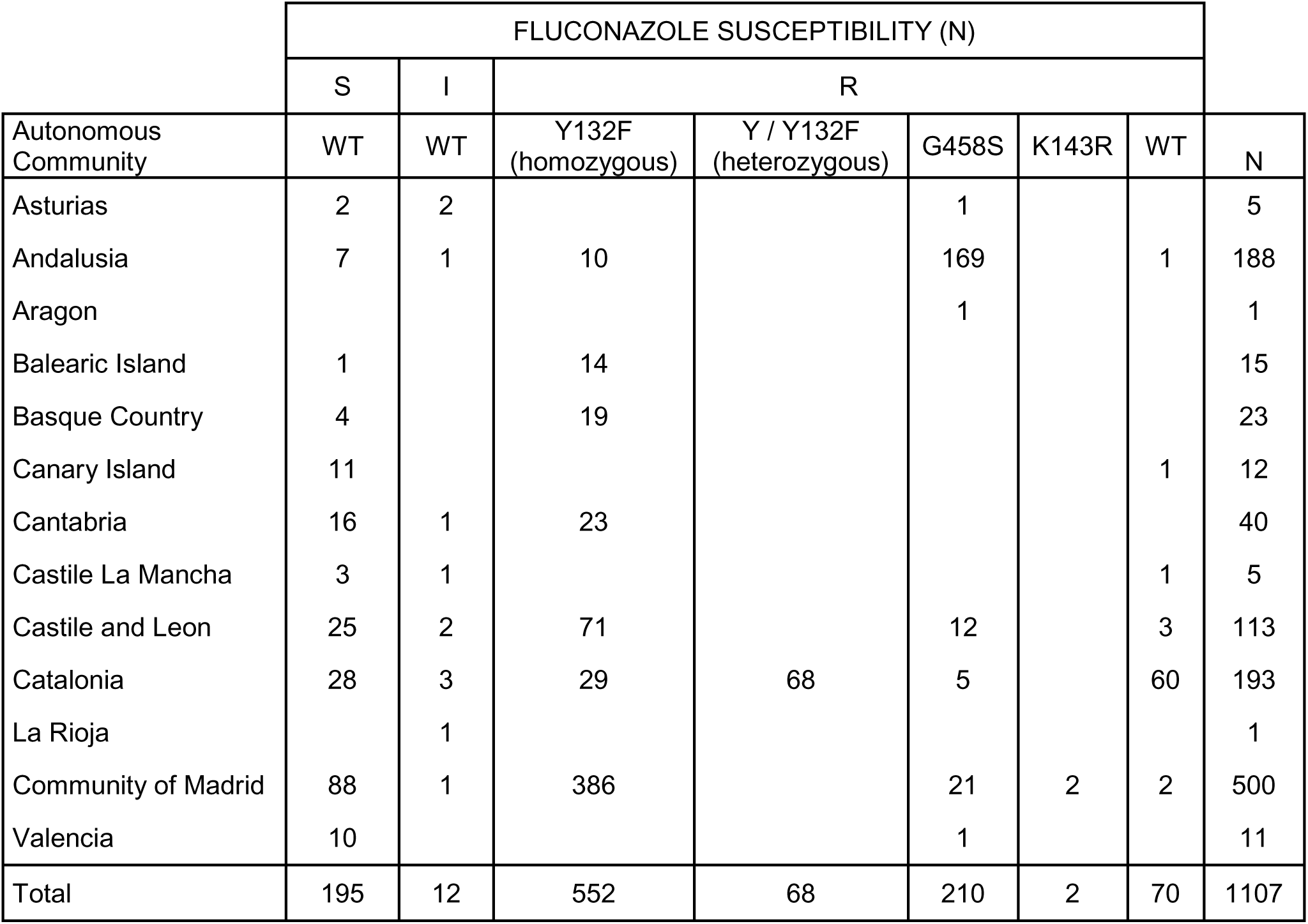
Distribution of fluconazole susceptible and resistant strains in different geographical areas from Spain.

**Table 3:** Distribution of type of samples from where the yeast strains were isolated. S, susceptible, I, intermediate (susceptible increase dose), R, resistant.

| Type and source of sample |  | FLZ susceptibility |  |  | TOTAL |
| --- | --- | --- | --- | --- | --- |
|  |  | S | I | R |  |
| Colonization screening | Anal exudate | 1 |  | 60 | 61 |
|  | Skin | 31 | 3 | 340 | 374 |
|  | Oropharyngeal exudate | 7 |  | 11 | 18 |
| Superficial | Sputum | 2 |  | 4 | 6 |
|  | Otic exudate | 1 |  |  | 1 |
|  | Ear exudate | 7 |  | 3 | 10 |
|  | Wound | 15 |  | 26 | 41 |
|  | Vaginal exudate | 2 |  | 3 | 5 |
| Invasive | Abdominal source | 7 | 1 | 18 | 26 |
|  | Abscess | 6 |  | 7 | 13 |
|  | Blood | 87 | 2 | 322 | 411 |
|  | Bone | 4 |  |  | 4 |
|  | CSF |  |  | 6 | 6 |
|  | Heart | 1 |  | 2 | 3 |
|  | Lung Biopsy | 2 |  | 1 | 3 |
|  | Muscle biopsy | 1 |  | 1 | 2 |
|  | Respiratory | 8 | 2 | 55 | 65 |
| Urine |  | 9 | 3 | 27 | 39 |
| Environmental |  | 1 |  | 12 | 13 |
| Unknown |  | 3 | 1 | 2 | 6 |
| TOTAL |  | 195 | 12 | 900 | 1107 |

Most infection-related strains were isolated from blood cultures (70%), followed by respiratory samples (13%), including bronchoalveolar lavages (BALs). Notably, a considerable proportion of isolates originated from non-invasive sites and were obtained through colonization screening.

### Antifungal susceptibility

Fluconazole resistance was observed in 81% of strains (900/1,107), with 1.1% classified as intermediate (susceptible increased exposure, 12/1,107). Resistance rates were 24.6% for itraconazole (273/1,107), 68.3% for voriconazole (757/1,107), and 24.2% for posaconazole (268/1,107), with an additional 12.5% showing intermediate susceptibility to voriconazole (139/1,107). Minimum inhibitory concentrations (MICs) for isavuconazole among fluconazole-resistant strains ranged from 0.016 to 8 mg/L. All fluconazole-resistant isolates remained fully susceptible to amphotericin B and echinocandins (data not shown).

### Microsatellite-based genotyping

Microsatellite genotyping of 1,067 strains was performed to assess genetic relatedness and comparison with previously described clones [31]. Higher genotypic diversity was observed among fluconazole-susceptible strains, whereas most resistant isolates clustered within known clonal complexes or formed new genotypes (**Figure 2**). Resistant strains mainly grouped into four major clonal complexes (96, 10, 67, and 54). Within complexes 96 and 67, closely related genotypes differing at a single microsatellite locus were identified, suggesting microevolution within these lineages (**Figure 2**).

**Figure 2:**
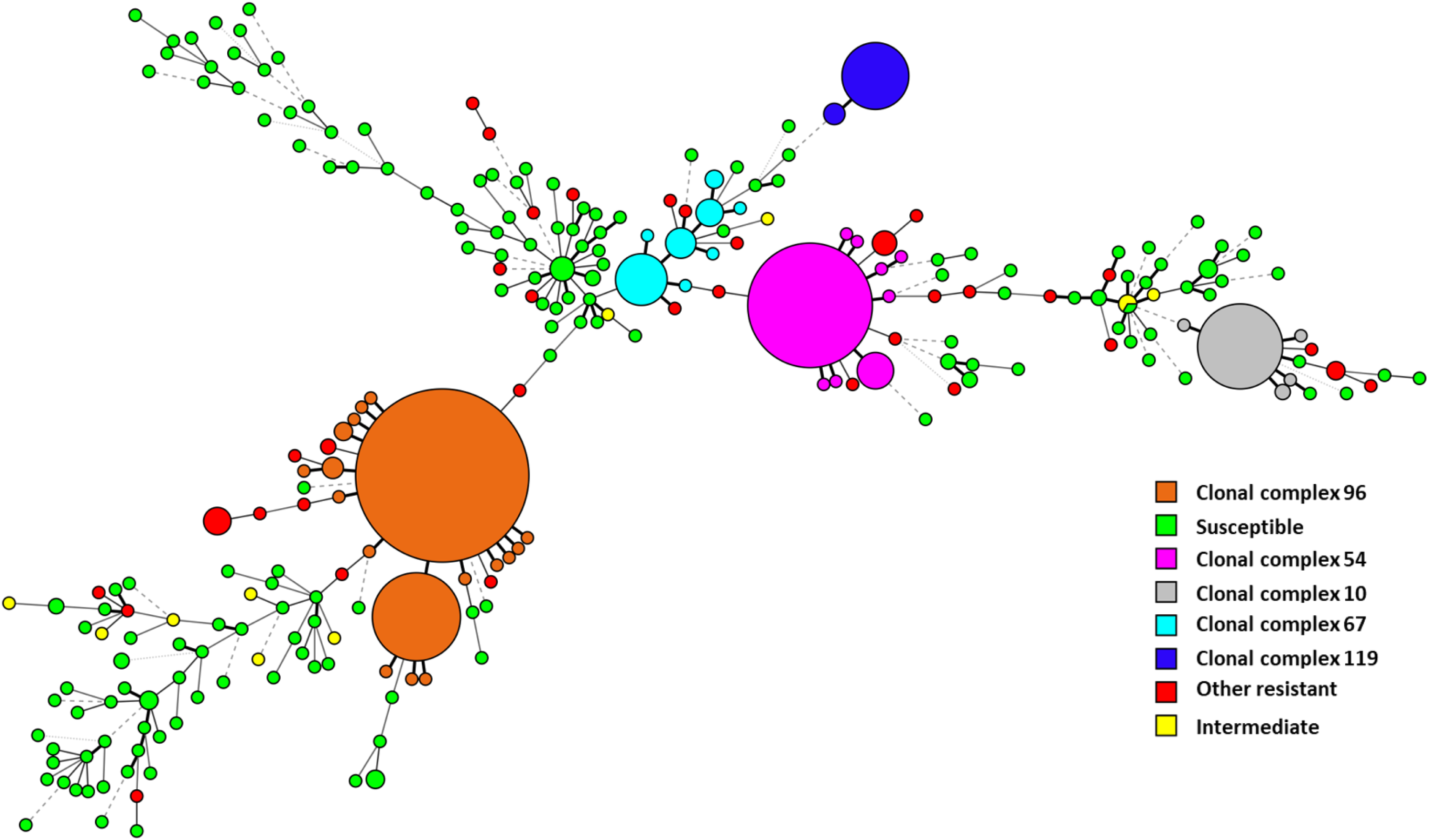
Minimum spanning tree representing the genetic relationships among the *C. parapsilosis* strains genotyped in this study. Each node corresponds to an individual strain, with distances reflecting genetic proximity. Clonal complexes, as defined in the Materials and Methods section, are highlighted using distinct colors: orange indicates clonal complex 96; pink, clonal complex 54; grey, clonal complex 10; light blue, clonal complex 67; and dark blue, clonal complex 119. Other Resistant strains are shown in red, isolates with intermediate susceptibility in yellow, and susceptible strains in green.

We identified distinct clusters associated with geographically close regions or hospitals **(Figure 3)**. Among them, the clones initially detected in the central region of Spain, first isolated in Madrid and Burgos (clone 96, according to [31]), was also found in other cities from Central Spain, such as Valladolid and Salamanca. Interestingly, this clone was also found in other distant cities in the south of Spain, such as Málaga.

**Figure 3:**
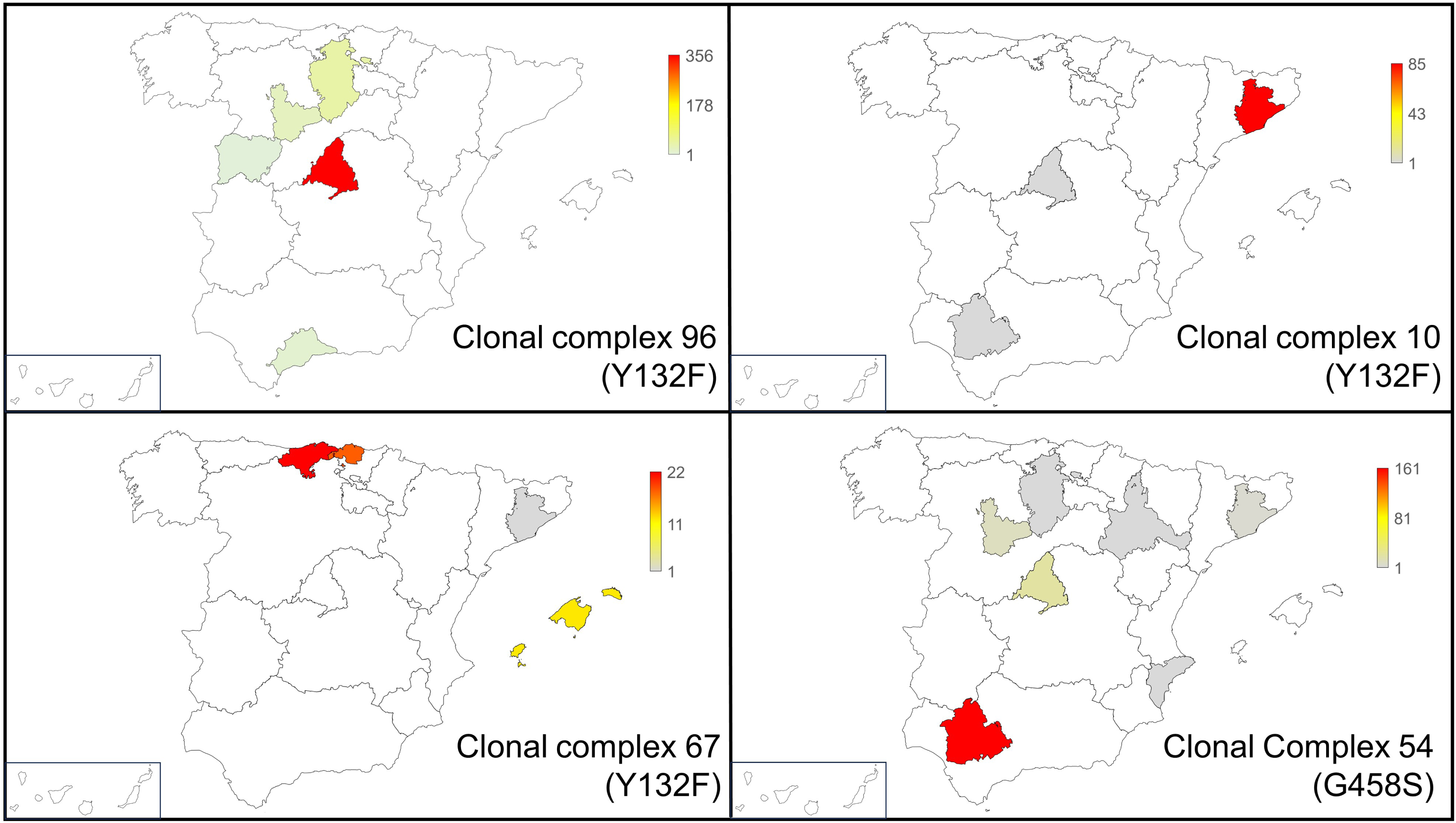
Geographical distribution of the main resistant genotypes harboring *ERG11* mutations. Each panel represents a distinct genotype (clonal complex), illustrating its spatial distribution across Spain: (a) clonal complex 96, (b) clonal complex 10, (c) clonal complex 67, and (d) clonal complex 54. Provinces are colored according to the number of resistant isolates, using a color scale ranging from grey (minimum value) to red (maximum value), with intermediate values represented by a gradient.

In Catalonia, two major clones were identified across multiple hospitals. Most strains belonged to genotype 10, previously described in this region, with random detection in Madrid and Andalusia (one strain each). Additionally, a distinct cluster of fluconazole-resistant strains (genotype 119) was identified at Sant Joan de Déu Hospital.

Furthermore, the clonal complex previously reported as circulating in the Balearic Islands and Cantabria (clone 67; [31]) has now also been detected in a hospital in the Basque Country and one isolate in Catalonia too.

Finally, in addition to these major clonal complexes, we identified a new large cluster of strains (clone 54) that first emerged in Andalusia, but was also detected in Valladolid, and at a lower prevalence in Madrid, Burgos, Zaragoza, Catalonia and Alicante.

### Azole resistance linked to *ERG11* mutations

In our previous study, we described three predominant genetic clones harboring the Erg11^Y132F^ mutation [31]. Since that time, these clones have continued to be isolated from the majority of previously affected hospitals and have demonstrated further geographic spread, with detection now reported in additional healthcare facilities.

Sequencing of the *ERG11* gene revealed that, since 2022, a significant number of strains have harbored the Y132F or G458S mutations **(Table 4).** Most G458S-carrying isolates clustered in the clonal complex 54. In total, strains carrying the G458S mutation have been isolated from 8 provinces and 13 different hospitals. Of note, we identified two strains harboring the G458S mutation that belonged to related, but slighted different genotypes (genotypes 55.2 from Catalonia and 55.4 from Asturias).

**Table 4.**
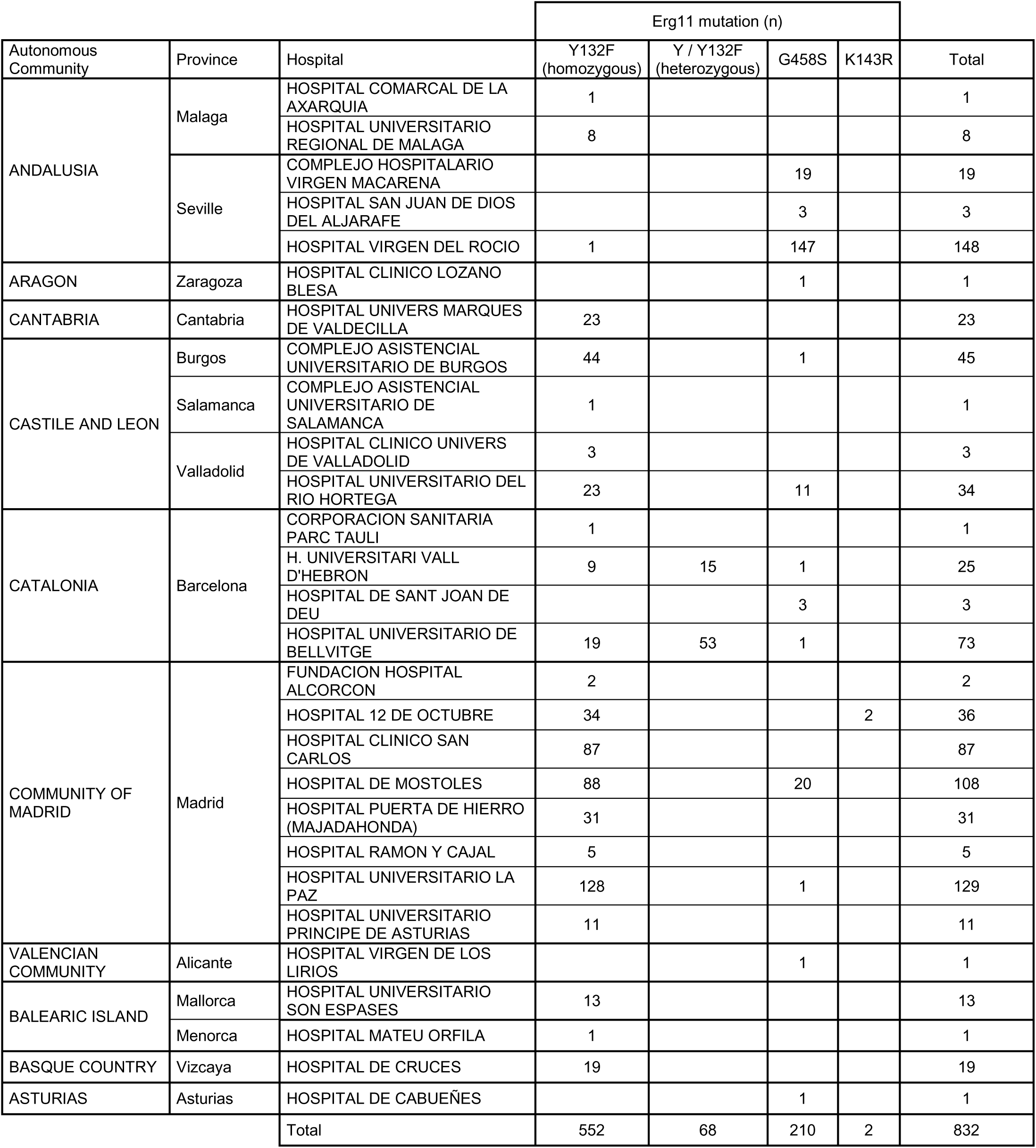
Distribution of the incidence of strains harboring Y132F, G458S and K143R mutations at Erg11 from different hospitals in Spain.

We also identified, although at lower prevalence, other resistant strains carrying alternative Erg11 mutations (specifically K143R) or exhibiting no detectable alterations in the *ERG11* gene. A summary of clonal complexes, resistance mechanisms and their geographical distribution is represented in **Figure 3**.

Cross resistance to other azoles vary depending on *ERG11* mutation (Table 5). All Y132F strains were fluconazole-resistant, with 80% also resistant to voriconazole (remaining intermediate) and low resistance to itraconazole (6.4%) and posaconazole (7.7%). In contrast, G458S strains showed high resistance to all azoles, including voriconazole (99.5%), itraconazole (80.9%), and posaconazole (74.2%).

**Table 5:** Cross resistance between azole antifungals in strains harboring the Y132F and G458S mutation.

|  | % Resistance |  |  |  |  |
| --- | --- | --- | --- | --- | --- |
|  | FLC | VOR |  | ITRA | POSA |
| Mutation | R | R | I | R | R |
| Y132F | 100 | 80 | 20 | 6.4 | 7.7 |
| G458S | 100 | 99.5 | 0.5 | 80.9 | 74.2 |

### Mutations in other genes involved in azole resistance

We also examined if the main clones harbouring mutations at *ERG11* had mutations at other genes that have been related to resistance to azoles by re-analyzing the previously-generated whole-genome sequencing data [47]. All the strains had the L643S substitution at Cdr4 and S208V and S304G at Erg6, suggesting that these changes represent in fact polymorphisms of the CDC317 reference strain. Strains from clonal complex 96 also harbored the F430I substitution at Mrr1. Of note, all the strains from clone 54 that had the G458S mutation also presented the G1330A mutation in the *TAC1* gene, which resulted in the D444N mutation in the Tac1 transcription factor. Furthermore, these strains also presented another significant genomic rearrangement, which involved a duplication of around 500,000 bp of chromosome 3 (NW_023503279.1, **Figure 4A)**. Interestingly, this duplicated region contained both the *ERG11* and *TAC1* genes.

**Figure 4:**
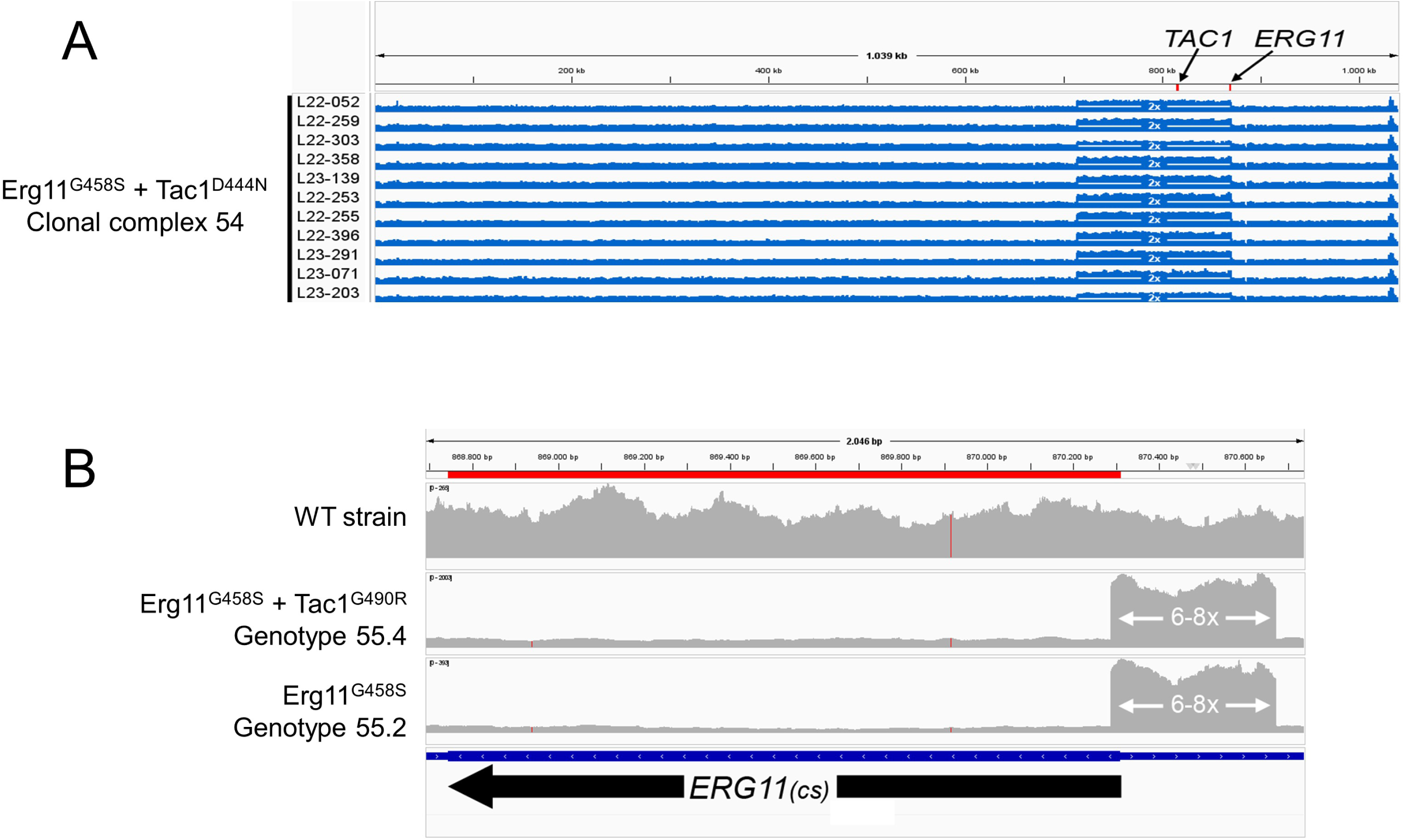
Genomic duplication and increase in copy number in strains harboring the G458S mutation at Erg11. A) Genomic duplication of the chromosome containing the *ERG11* and *TAC1* in strains from genotype 54 (Erg11^G458S^ and Tac1^D444N^). B) Genomic coverage of *ERG11* gene in strains from genotypes 55.2 and 55.4. As annotated in the CDC317 genome, *ERG11* gene is located in the minus strain, as indicated by the arrow below the gene.

The strain from genotype 55.2 (G458S) had a different mutation at the *TAC1* gene (G1468C), which caused the G490R mutation at the Tac1 transcription factor. In contrast, the strain from genotype 55.4 that also had the G458S mutation did not present any mutation at the *TAC1* gene. While these two did not present the duplication present in the rest of the strains with this mutation at the *ERG11* gene, they both showed an increased number of copies (6-8x) of a small region of 385 bp that partially affected the promoter and beginning of the coding region of the *ERG11* gene (363 bp from the promoter and first 22 bp from the coding region, **Figure 4B**).

We found that one strain that harbored the K143R mutation at Erg11 (strain L24/020) also had an increase in the number of copies of the *ERG11* gene **(Figure 5).** Regarding other resistant strains without *ERG11* alterations, we found that they presented mutations in transcription factors such as Mrr1 and Upc2 **(Table 6)**. Two of these resistant strains also had the G220N substitution at Erg6.

**Figure 5:**
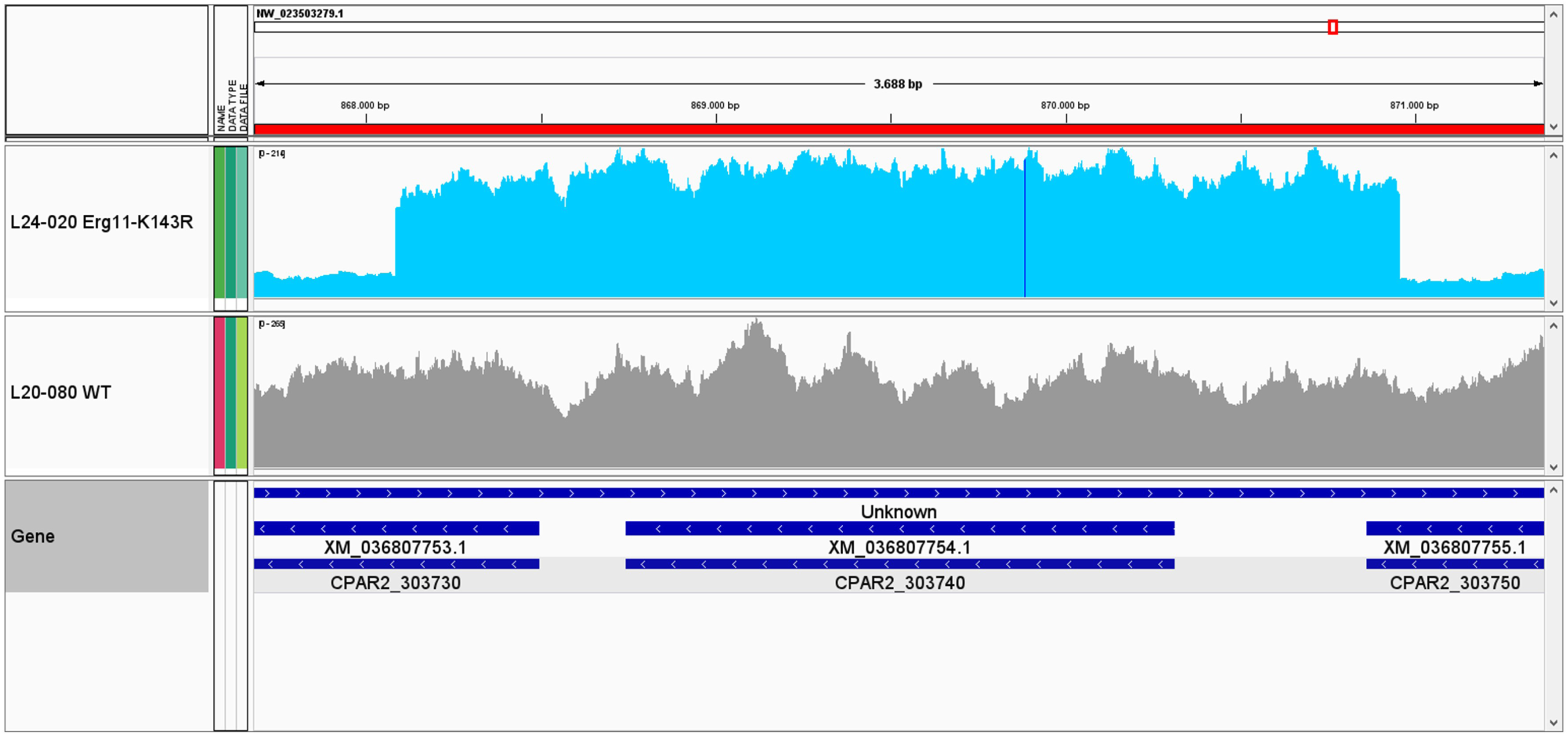
Coverage of *ERG11* reads in strain L24-020 harboring the K143R substitution in *ERG11*. The upper panel shows sequencing read coverage for strain L24-020 (*ERG11* K143R), exhibiting an approximately 7 increase in copy number, while the lower panel shows a wild-type strain (L20-080).

**Table 6.** Variations and mutations observed in Fluconazole resistant strains with or without mutations at *ERG11*.

|  | Mrr1<br>CPAR2_807270 | Upc2<br>CPAR2_207280 | Tac1<br>CPAR2_303510 | Cdr1<br>CPAR2_405290 | Cdr4<br>CPAR2_600730 | Erg6<br>CPAR2_405010 | Erg25<br>CPAR2_801410 | Erg11<br>CPAR2_303740 |
| --- | --- | --- | --- | --- | --- | --- | --- | --- |
| L22/299 | P295S<br>(heterozygous) | - | - | I1287V | L643S | S208V<br>S304G | D9N (het) | - |
| L23/089 | K177N | - | - | - | L643S | S208V<br>G220N<br>S304G | - | - |
| L23/346 | K177N<br>G483Q (het) | - | - | - | L643S | S208V<br>G220N<br>S304G | I307T | - |
| L25/219 | G483Q (het) | N455D | - | I969T<br>I1287V | L643S | S208V<br>S304G | - |  |
| GENOTYPE<br>96 | F430I | - | - | - | L643S<br>T1180I | S208V<br>S304G | - | Y132F |
| GENOTYPE<br>10 | - | - | - | - | L643S<br>T764I | S208V<br>S304G | - | Y132F |
| GENOTYPE<br>15 | - | - | - | - | L643S<br>T764I (het) | S208V<br>S304G | - | - |
| GENOTYPE<br>67 | - | - | - | - | L643S | S208V (het)<br>S304G (het) | - | Y132F |
| GENOTYPE<br>54 | - | - | D444N | - | L643S | S208V<br>S304G | - | G458S |
| L23/273<br>(GENOTYPE 55.2) | - | - | D490R | - | L643S | S208V (het)<br>S304G (het) | - | G458S |
| L23/087<br>(GENOTYPE 55.4) | - | - | - | - | L643S | S208V<br>S304G | - | G458S |

### Determination of *ERG11* copy number

To confirm that the strains from clonal complex 54 (Erg11^G458S^ + Tac1^D444N^) had a duplication of the *ERG11* gene, we developed a real-time PCR to compare the number of copies of *ERG11* related to *ACT1* (unicopy gene, see M&M). As shown in **Table 7**, the number of copies of the *ERG11* gene in strains from clone 54 was close to 2. No increase in copy number of the strains from genotypes 55.2 and 55.4 was observed (data not shown).

**Table 7:** *ERG11* copy number estimated by real time PCR.

|  | Strain | *ERG11/ACT1*<br>copy number | Average ± s.d. |
| --- | --- | --- | --- |
| Susceptible<br>strains | L23/221 | 0.98 | 0.89 ± 0.27 |
|  | L23/222 | 0.62 |  |
|  | L23/223 | 0.99 |  |
|  | L23/224 | 0.68 |  |
|  | L23/225 | 0.75 |  |
|  | L23/226 | 1.34 |  |
| G458S strains<br>(genotype 54) | L23/071 | 2.00 | 2.11 ± 0.35 |
|  | L23/139 | 2.18 |  |
|  | L23/203 | 2.08 |  |
|  | L23/216 | 1.68 |  |
|  | L23/291 | 2.64 |  |

### Correlation between Erg11 mutations, genotype and antifungal susceptibility profiles

All the fluconazole susceptible and resistant strains remained fully susceptible to other antifungals, such as AmB **(figure 6A)** and echinocandins (data not shown), although strains harbouring the Y132F mutation presented higher MIC values (1-fold dilution higher) than susceptible and G458S strains. Regarding FLZ susceptibility, strains from genotype 10 had lower MIC values than those from clones 96 and 67. This was mainly due to the fact that this clonal complex included both Y132F mutants in homozygosis and heterozygosis. When we analysed homozygous and heterozygous Y132F strains from clone 10, we observed that the heterozygous strains had lower MIC values that the homozygous strains **(Figure 6B),** which is in agreement with previous findings [31]. Although strains from clone 96 harboured the Y132F at Erg11 and F430I substitution at Mrr1, they did not show higher MIC values to other strains with Y132F mutations, suggesting that the F430I mutation at Mrr1 did not have a significant impact on fluconazole resistance **(Figure 6).** For strains with the Y132F mutation, we found that there was a slight increase in the MIC distribution to ITZ and POS, but most of them were still categorized as susceptible according to the established breakpoints.

**Figure 6:**
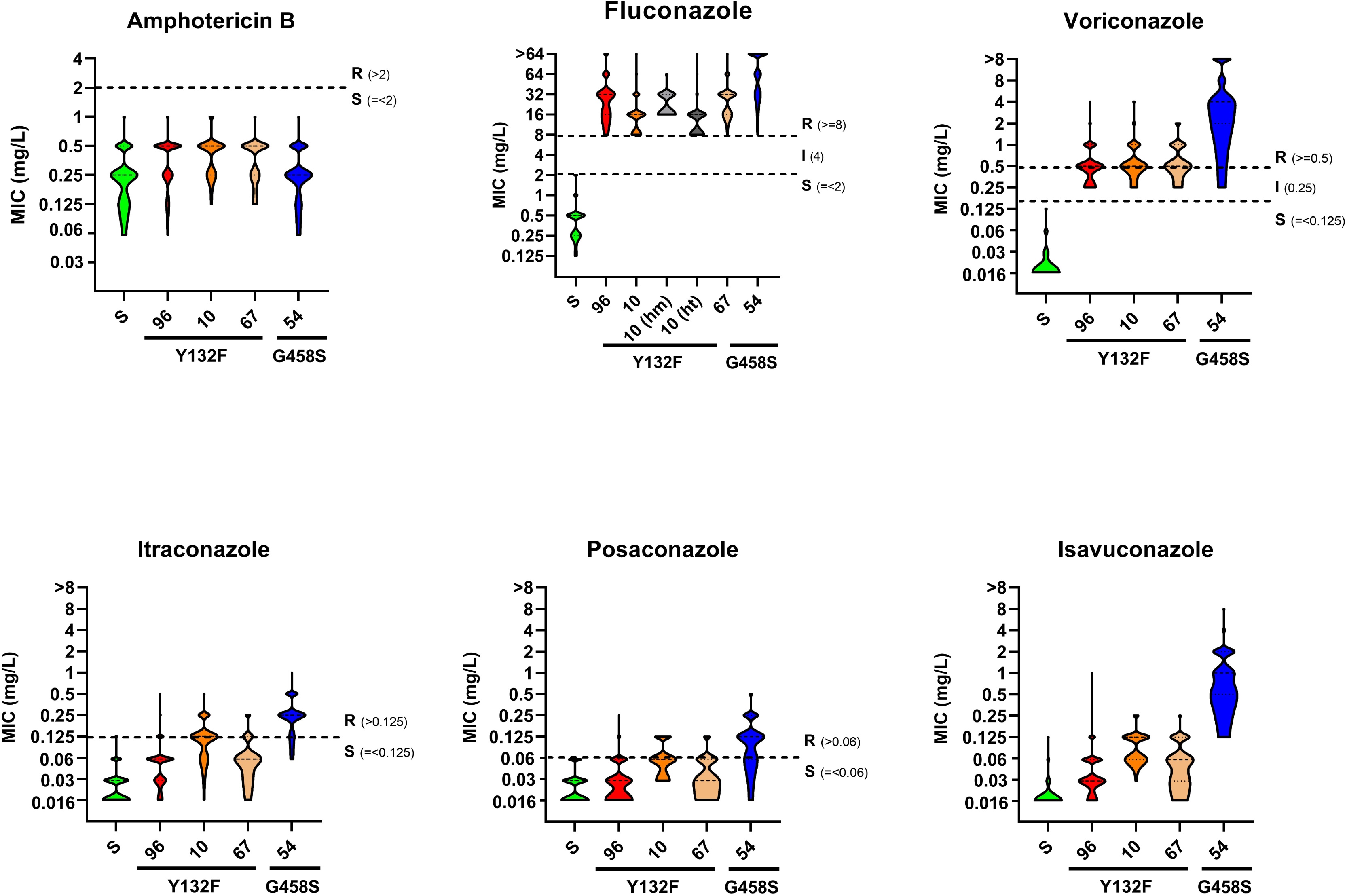
Distribution of MIC values for azoles and amphotericin B in strains harboring the Y132F and G458S mutations, as well as in susceptible strains. Violin plots show the distribution of MIC values across clonal complexes. Within clonal complex 10 in the fluconazole panel, strains are classified as heterozygous (hm) or homozygous (ht) for the mutation. Horizontal dashed lines indicate clinical breakpoints for susceptible (S), intermediate (I), and resistant (R) categories, where applicable.

Isolates with Erg11^G458S^ + Tac1^D444N^ (clone 54) had significantly higher MICs to FLZ, VOR and in particular to ISV than strains harbouring the Y132F mutation **(Figure 6B, C and F**, p value < 0.05 for all comparisons). G458S strains from complex 54 had a moderate increase in the MIC values to ITZ and POS compared to the Y132F substitution, and most of the isolates presented MIC values above the defined breakpoints and were categorized as resistant **(Figure 6D and E)**.

We also compared the antifungal susceptibility of the two strains from genotypes 55.2 (Erg11^G458S^ + Tac1^D490N^) and 55.4 (Erg11^G458S^). Strikingly, the strain harbouring the Erg11^G458S^ but no mutations at the Tac1 transcription factor showed lower resistance to azoles that other Erg11^G458S^ strains carrying a Tac1 mutation **(Table 8).**

**Table 8.** Antifungal susceptibility of strains harboring the G458S mutation at Erg11.

|  |  | AmB | 5FC | FLZ | ITRA | VOR | POS | ISA | CASPO | MICA | ANIDU |
| --- | --- | --- | --- | --- | --- | --- | --- | --- | --- | --- | --- |
| Erg11<sup>G458S</sup><br>+<br>Tac1<sup>D444N</sup><br>Clone 54 | Min | 0.06 | 0.125 | 16 | 0.06 | 0.5 | 0.03 | 0.125 | 1 | 0.5 | 0.25 |
|  | Max | 1 | 0.25 | >64 | 0.5 | >8 | 0.5 | 16 | 2 | 1 | 2 |
|  | GM | 0.3 | 0.1 | >64 | 0.3 | 4.9 | 0.1 | 0.9 | 1.2 | 0.7 | 1.1 |
|  | Mode | 0.25 | 0.125 | >64 | 0.25 | >8 | 0.125 | 2 | 1 | 1 | 1 |
| Erg11<sup>G458S</sup><br>+<br>Tac1<sup>G490R</sup><br>Genotype 55.2 | MIC | 0.5 | 0.125 | >64 | 0.5 | 4 | 0.25 | 2 | 1 | 0.5 | 1 |
| Erg11<sup>G458S</sup><br>Genotype 55.4 | MIC<br>(Two repetitions) | 0.06<br>0.25 | 0.125<br>0.125 | 8<br>8 | 0.25<br>0.25 | 0.25<br>0.5 | 0.125<br>0.125 | 0.06<br>0.25 | 2<br>2 | 1<br>1 | 0.5<br>1 |

### Phylogenetic analysis by whole genome sequencing

We performed a phylogenetic analysis using the complete genomes of representative strains from the different clones obtained from different hospitals, and also susceptible isolates from our collection. The genome wide analysis also included a selection of strains isolated in previous years (from 2019 to 2022) which covered strains from different hospitals and from the main genotypes and clonal complexes already defined (31). Phylogenetic analyses of whole genomes confirmed the main clonal clusters obtained by microsatelites. As shown in **Figure 7**, the main clones harboring the Y132F mutations and G458S clustered in the unique clades. Interestingly, resistant strains with the Erg11^Y132F^ were clonally present in some hospitals since 2019, confirming the local transmission in these centers. In our analysis based on clustering of the whole genomes, we could not differentiate the micro-evolved genotypes found in the microsatellite analysis within the same clonal complex, supporting the notion that clonal complex include genotypes that differ in one microsatellite [45]. In a complementary study [47], we have shown that these strains constitute new clones from new subclades that do not cluster with genomes from other published outbreaks. In particular, some of these clones have been defined as new subclades (clone 10 as subclade 4.1, clone 67 as subclade 4.2 and clone 54 as subclade 4.3). Interestingly, we identified several susceptible strains with no mutations at *ERG11* (defined as genotype 15 by microsatellite analysis) that clustered together with the resistant strains from clonal complex 10, suggesting that these Y132F resistant strains might have been originated from this genotype 15. Subclade 4.3 included all the strains harboring the Erg11^G458S^ mutation. Strains from clone 96 belonged to the previously defined clade 5. Notably, two isolates carrying the G458S mutation (strain L22-087 from genotype 55.2, and L23-273 from 55.4) were genetically distinct from the main cluster, although they still grouped within the same clade.

**Figure 7:**
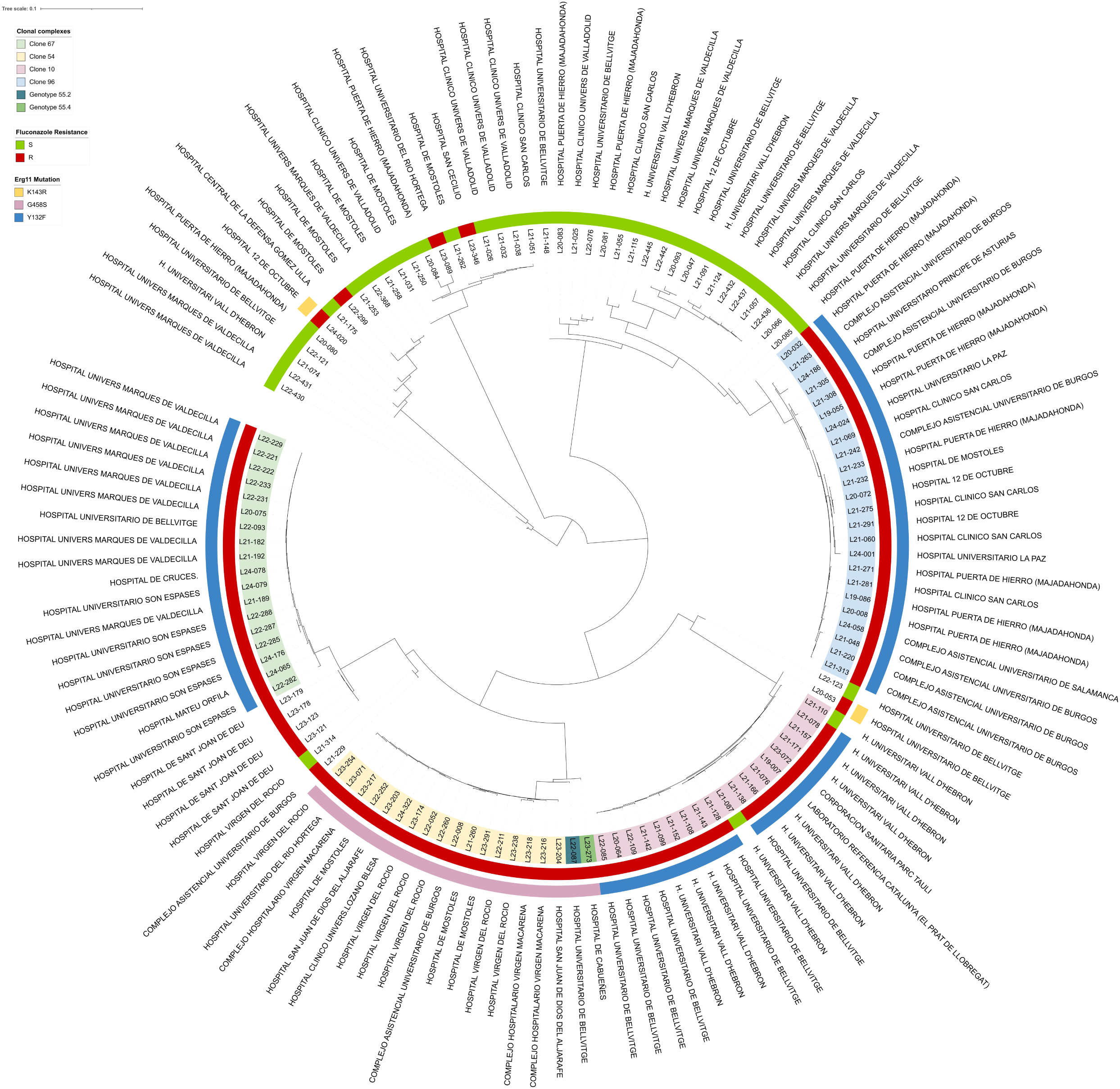
Phylogenetic tree based on whole-genome sequencing of susceptible strains and representative fluconazole-resistant isolates, including wild-type (WT) susceptible strains, strains harboring the Y132F, G458S, and K143R mutations, as well as resistant strains lacking *ERG11* mutations. Isolates are annotated according to their hospital of origin. The legend summarizes clonal complexes, fluconazole susceptibility, and mutation profiles. Isolates are annotated according to their hospital of origin.

## Discussion

The recent worldwide rise of fluconazole resistance in *C. parapsilosis* poses a new challenge in antifungal resistance [8, 55]. In Spain, this phenomenon has been particularly noticeable since 2020 in numerous hospitals, coinciding with the onset of the COVID-19 pandemic. These studies described the different clonal complexes carrying the Y132F mutation at Erg11. Some of these clones have been associated with specific geographical regions [9, 17, 31]; however, the same clone has also been detected in geographically distant areas [31].

To date, the prevalence of isolates carrying the G458S mutation has been limited compared with the outbreaks caused by strains harboring the Y132F mutation [17, 19, 31]. In this study, we report the isolation of a considerable number of *C. parapsilosis* isolates carrying the Erg11^G458S^ mutation from hospitals located in different geographical regions. This pattern contrasts with the higher genetic diversity observed among *C. parapsilosis* strains harboring the Y132F mutation in Spain. It remains unclear whether the emergence of the G458S mutation is restricted to one or a limited number of genotypes due to the relatively small number of such isolates analyzed worldwide compared with Y132F-carrying strains.

One particularly relevant aspect of this work is the origin of these isolates. Most were detected through surveillance screening of non invasive samples, mainly from skin, suggesting the existence of potential sources of patient colonization or infection in specific hospital settings. Since antifungal susceptibility testing of isolates from non-invasive sources is not routinely performed in many hospitals, this practice gap may contribute to the unnoticed spread and dissemination of specific resistant clones. Our findings support the recommendation to periodically perform antifungal susceptibility testing on a selected number of *C. parapsilosis* isolates from non-invasive sources to ensure early detection of transmission events. This could significantly contribute to prevent the establishment of clinical outbreaks by implementing preventive measures such as targeted cleaning and decontamination of hospital areas at risk of becoming reservoirs of infection.

Most outbreaks of *C. parapsilosis* recently reported in Spain have been caused by clonal resistant isolates, suggesting that these strains may display phenotypic traits that confer a selective advantage for dissemination. Several studies have not found an association between the emergence of these outbreaks and the use of triazoles in the affected centers [15, 24, 31]. In this context, different studies have addressed whether fluconazole resistant strains present different phenotypic features that could explain their capacity to establish outbreaks. Strikingly, it has been shown that these clones have a diminished capacity to form biofilms and induce pseudohyphae [28, 56]. Consequently, we argue that selection of resistant isolates is likely driven by multiple factors, including environmental conditions, patient susceptibility to colonization, infection prevention and control practices, and intrinsic microbiological characteristics.

When analyzing the antifungal susceptibility profiles of isolates carrying the two main mutations, we observed that the G458S substitution had a greater impact on the resistance phenotype. In particular, isolates harboring the G458S substitution alone exhibited higher levels of resistance to all azoles tested, resulting in the vast majority being classified as resistant to fluconazole, voriconazole, itraconazole, and posaconazole. This resistance profile contrasts with that reported for isolates carrying the Y132F mutation, which are typically resistant only to fluconazole and voriconazole.

A similar pattern has been described in a study investigating a prolonged outbreak of fluconazole-resistant *C. parapsilosis* initially detected in multiple healthcare centers in Berlin, Germany [36]. Interestingly, in that study, they identified strains with the G458S that cluster with the strains with the same mutation from this study [47].

At present, the molecular basis underlying the differences in antifungal susceptibility between isolates harboring these two mutations remains unknown. Overall, our data are consistent with the observation that both mutations primarily affect susceptibility to short-chain triazoles (voriconazole, fluconazole, and isavuconazole), while their impact appears to be more limited for azoles with extended side chains, such as posaconazole and itraconazole. Evidence from *Candida albicans* indicates that different amino acid substitutions contribute to varying levels of antifungal resistance. From a structural perspective, the Y132 aminoacid substitution is located within the active site of Erg11 and it interacts with the triazole ring of azoles, contributing to the hydrogen-bonding network that stabilizes heme iron coordination. Substitution of tyrosine (T) with phenylalanine (F) removes the hydroxyl group, thereby weakening critical polar interactions and reducing enzyme affinity for short-chain azoles [57–59]. In contrast, itraconazole and posaconazole are less affected by this substitution, likely because their lipophilic side chains interact with additional residues along the ligand-access channel of the enzyme, partially compensating for the reduced binding affinity.

Differently to Y132F, the G458S substitution, located in the C-terminal region of Erg11p near the transmembrane domain, is predicted to influence local protein folding or membrane orientation rather than directly affecting the active site. However a striking finding is that most G458St strains also carry a Tac1 transcription factor mutation and a ∼500 kb duplication on chromosome 3 affecting *ERG11* and *TAC1*, consistent with previously reported mechanisms involving gene duplications [60]. These additional alterations likely contribute to the higher resistance observed. This is supported by one strain with the G458S substitution but no Tac1 mutation, which showed the lowest resistance. We also identified a strain with increased *ERG11* copy number, a change reported in other strains [61].

Overall, our results indicate that fluconazole resistance arises from multiple coexisting mechanisms, including genomic duplications, increased target gene copy number, and mutations in several genes.

Supporting this idea, in the complementary work by Schikora-Tamarit et al [47] which describes an extensive genome-wide association study (GWAS) including many of the strains described in this manuscript, it was found that resistance to fluconazole cannot be explained by a single underlying mechanism. Additionally, that study identified new genomic variations associated with previously described phenotypic traits involved in virulence [56] and with the clinical sample from which the strains were isolated. Together, these two studies open new perspectives to understand the factors that determine the emergence of fluconazole resistance in *C. parapsilosis*.

In conclusion, although recent reports have described outbreaks of FNS *C. parapsilosis*, this study provides one of the largest collections of isolates with the G458S mutation and also reports the expansion of resistant clones carrying Y132F. The presence of these resistant isolates in colonization screenings suggests that these isolates may become stable components of the human microbiota, acting as reservoirs for future infections with antifungal treatment failure. Therefore, we advocate for routine surveillance programs in clinical practice, including periodic antifungal susceptibility testing, to enable early detection and prevention of outbreaks. We also recommend genetic characterization of resistant isolates to better define their epidemiology and limit their spread across healthcare centers.

## Acknowledgements and Funding

This work has been partially supported by Gilead S.L. O. Z. and L.A-F are funded by grant AESI-2024 PI24CIII/00051 (Instituto de Salud Carlos III). O.Z. is also funded by grant PID2023-148686OB-I00 by MICIU/AEI/10.13039/501100011033 and by FEDER, UE (Ministerio de Ciencia, Innovación y Universidades). Elena López Peralta is funded by a contract by the Comunidad Autónoma de Madrid (AYUDAS PARA LA CONTRATACIÓN DE PERSONAL INVESTIGADOR PREDOCTORAL EN FORMACIÓN, PIPF-2022/SAL-GL-24953). T.G. acknowledges support from the Spanish Ministry of Science and Innovation, grant number PID2021-126067NB-I00 founded by MICIU/AEI /10.13039/501100011033 and by FEDER, as well as well as support from “La Caixa” foundation (grant number LCF/PR/HR21/00737, CI23-20260). MAST acknowledges his AI4S fellowship within the “Generación D” initiative by Red.es, Ministerio para la Transformación Digital y de la Función Pública, for talent attraction (C005/24-ED CV1), funded by NextGenerationEU through PRTR.

